# A Scalable Toolkit for Modeling 3D Surface-based Brain Geometry

**DOI:** 10.1101/2025.08.06.668521

**Authors:** Yanghee Im, Leila Nabulsi, Melody J.Y. Kang, Sophia I. Thomopoulos, Ana M. Diaz Zuluaga, Anders M. Dale, Andriana Karuk, Annabella Di Giorgio, Benson Mwangi, Boris Gutman, Bronwyn Overs, Carlos López Jaramillo, Colm McDonald, Dan J. Stein, Dara M Cannon, David Glahn, Diego Hidalgo-Mazzei, Diliana Pecheva, Dominik Grotegerd, Edith Pomarol-Clotet, Eduard Vieta, Emilie Olie, Enric Vilajosana Chertó, Fabio Sambataro, Fleur Howells, Freda Scheffler, Geraldo Busatto, Gerard Anmella, Giovana B. Zunta-Soares, Gloria Roberts, Henk Temmingh, Ian Gotlib, Ingrid Agartz, Jair C. Soares, James A. Karantonis, James Prisciandaro, Janice M. Fullerton, Joaquim Radua, Jonathan Savitz, Josselin Houenou, Kang Sim, Kenichiro Harada, Klaus Berger, Koji Matsuo, Lakshmi Yatham, Lars Tjelta Westlye, Lisa Eyler, Lisa Furlong, Luisa Klahn, Marco Hermesdorf, Marcus V. Zanetti, Matthew Kempton, Matthew Sacchet, Mikael Landen, Mon-Ju Wu, Pedro Rosa, Philip Mitchell, Pravesh Parekh, Raymond Salvador, Rayus Kuplicki, Salvador Sarró, Susan Rossell, Tamsyn Van Rheenen, Theodore Satterthwaite, Tilo Kircher, Tomas Hajek, Udo Dannlowski, Xavier Caseras, Yuji Zhao, Ole A. Andreassen, Paul M. Thompson, Christopher R.K. Ching, the ENIGMA Bipolar Disorder Working Group

## Abstract

3D surface-based computational mapping is more sensitive to localized brain alterations in neurological, developmental and psychiatric conditions than traditional gross volumetric analysis, providing fine-scale 3D maps of a wide range of surface-based features. Here we introduce a scalable toolkit for large-scale computational surface analysis, with efficient algorithms for multisite data integration, statistical harmonization, accelerated multivariate statistics, and visualization. We showcase the utility of the toolkit by mapping subcortical shape variations and factors that affect them across 21 international samples from the ENIGMA Bipolar Disorder Working Group (N=3,373).

## 1 Introduction

Alterations in gross subcortical volumes are a common signature across neurological [1,2] and psychiatric disorders (e.g., smaller hippocampal volumes in mm^3^) [3–5]. Surface-based 3D brain features offer enhanced spatial specificity to local (and sometimes complex) morphometric alterations (e.g., thickness or surface area) that are not captured when measuring gross volumetrics (e.g., whole hippocampal volume). Bipolar disorder (BD) is associated with brain alterations including smaller hippocampal and thalamic gross volumes [6]. Commonly prescribed treatments for BD are associated with differential patterns of brain morphometry, with lithium treatment associated with larger brain volumes, and antiepileptic and antipsychotic treatments associated with smaller volumes [3]. These treatment associations have been shown to influence machine learning diagnostic performance in BD [7] as well as other psychiatric disorders like obsessive compulsive disorder [8]. Finer-scale mapping of such treatment-related brain variations in large, multisite data could provide more clinically-relevant insights into predicting and monitoring treatment response.

The Enhancing Neuro Imaging Genetics through Meta-Analysis (ENIGMA) Consortium is a global effort uniting over 2,000 scientists to conduct the largest neuroimaging studies of over 30 brain diseases and disorders [9]. ENIGMA applies standardized neuroimaging processing and analysis protocols to existing data samples collected across the world to identify disease effects on brain structure and function. The ENIGMA Subcortical Shape Pipeline (https://github.com/ENIGMA-git) offers a standardized protocol for deriving local radial distance and surface Jacobian metrics along thousands of surface vertices from the hippocampus, amygdala, accumbens, caudate, putamen, pallidum and thalamus. The ENIGMA Subcortical Shape Pipeline has been used in the largest international analyses of subcortical anatomy in schizophrenia [10], major depression [11], substance use [12], Parkinson’s disease [2], 22q11.2 deletion/duplication syndrome [13]. Here, building on the ENIGMA Subcortical Shape Pipeline, we provide a Python-based workflow **(Fig. 1**) to support future projects performing multi-site harmonization, vertex-wise statistical modeling, machine learning analysis, and visualization of subcortical shape data. To address the computational challenges of high-dimensional vertex-wise data, our toolkit incorporates FEMA (Fast and Efficient Mixed-Effects Algorithm), a statistical framework to accelerate the computation of surface-based statistics [14]. In addition, the workflow integrates interpretable machine learning methods including Logit-TVL1, which combines sparsity and spatial smoothness constraints to classify and localize brain shape alterations [15, 16]. Comprehensive visualization tools are also provided to represent mass univariate statistical and machine learning results onto 3D subcortical surfaces, improving interpretability.

**Fig. 1.**
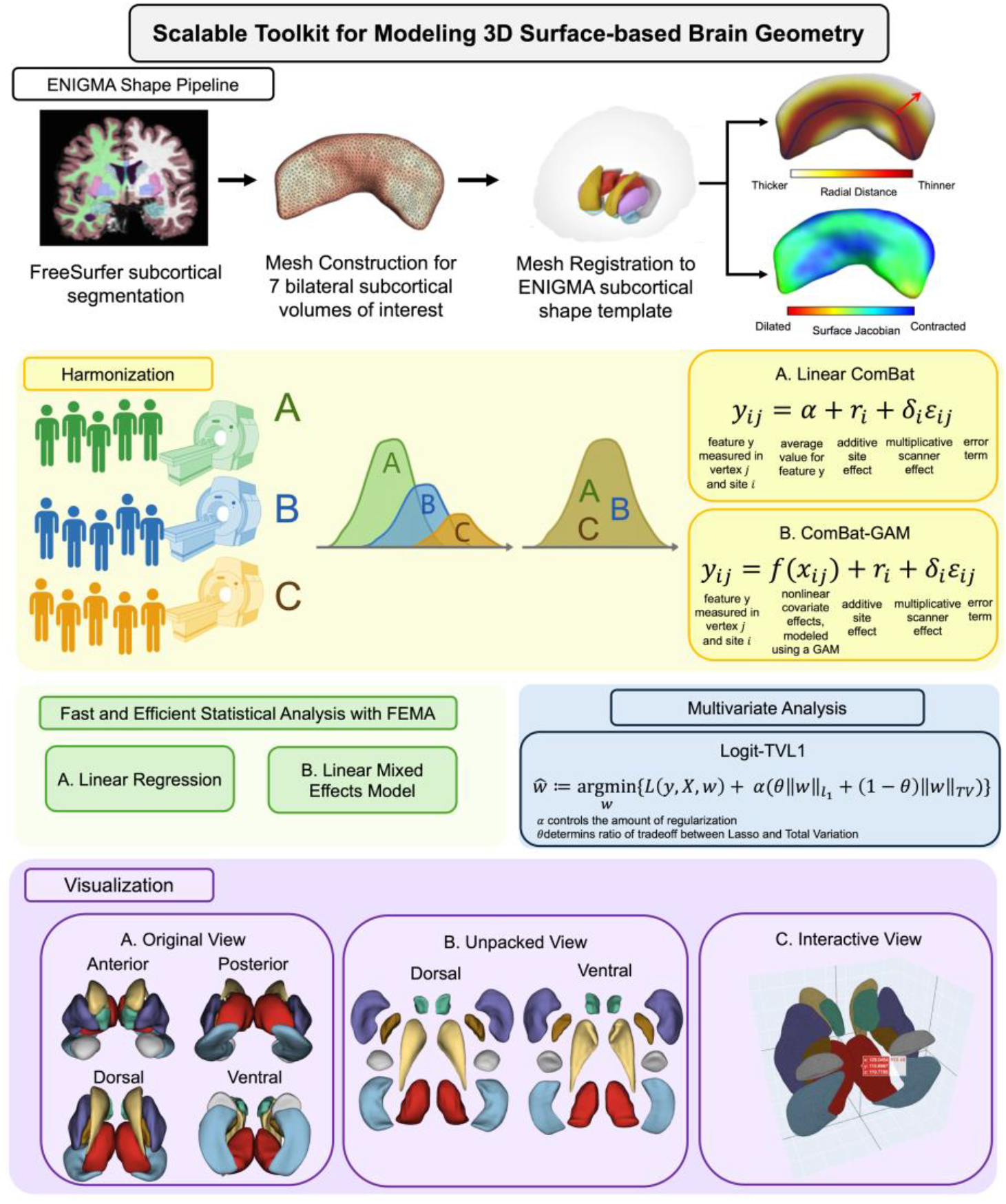
Overview of the proposed toolkit for modeling 3D surface-based brain geometry.

## 2 Methods

### 2.1 Data

3D T1-weighted brain MRI data from 21 independently collected study samples (BD=1,487, healthy controls (HC)=1,886; age: 8.0 - 83.0 years, 42% male) were segmented using the ENIGMA-standard FreeSurfer (v5.3) protocol [17] to derive 7 bilateral gross subcortical volumes (hippocampus, amygdala, caudate, putamen, pallidum, thalamus, and nucleus accumbens) for comparison to shape analysis metrics.

**Table 1.** Demographic characteristics for 21 ENIGMA-BD sites.

| **Sample** | **# of BD/HC (% Male)** | **Age Mean (SD)** |
| --- | --- | --- |
| Barcelona | 102/117 (46%) | 41.5 ($\pm 9.4$) |
| BiDirect | 42/40 (43%) | 51.0 ($\pm 7.5$) |
| Cardiff | 91/61 (33%) | 40.0 ($\pm 8.7$) |
| CHRM2 | 42/54 (49%) | 41.5 ($\pm 13.0$) |
| COGSBD | 59/31 (50%) | 37.1 ($\pm 11.2$) |
| Halifax | 62/52 (38%) | 38.0 ($\pm 14.7$) |
| Houston | 258/171 (43%) | 26.5 ($\pm 14.7$) |
| IDIBAPS | 25/57 (43%) | 37.5 ($\pm 13.6$) |
| Japan | 155/557 (36%) | 46.0 ($\pm 15.9$) |
| KCL | 23/22 (31%) | 42.1 ($\pm 14.4$) |
| Montpellier | 90/64 (40%) | 38.0 ($\pm 11.1$) |
| MUSC | 68/26 (49%) | 38.7 ($\pm 9.3$) |
| NUIG | 70/95 (56%) | 36.7 ($\pm 12.2$) |
| Paris | 36/55 (51%) | 34.5 ($\pm 13.3$) |
| Penn | 58/88 (45%) | 38.2 ($\pm 9.8$) |
| SBA | 30/43 (44%) | 34.0 ($\pm 10.5$) |
| Singapore | 43/44 (45%) | 23.4 ($\pm 3.9$) |
| Sydney | 58/99 (41%) | 35.0 ($\pm 11.1$) |
| Tulsa | 68/90 (29%) | 22.8 ($\pm 4.5$) |
| UBC | 64/42 (47%) | 50.4 ($\pm 13.2$) |
| UCSD | 43/78 (39%) | 37.8 ($\pm 11.6$) |
| **Total** | **1487/1886 (42%)** | **38.0 ($\pm 15.0$)** |

### 2.2 ENIGMA Subcortical Shape Features

The existing ENIGMA Subcortical Shape Pipeline (https://github.com/ENIGMA-git) uses FreeSurfer subcortical volumes to derive two shape features from seven bilateral subcortical models. Each structure is modeled as a triangular surface mesh, where the points on the surface define the vertices that make up the overall geometry of the 3D shape. Triangular surface meshes are registered to a common ENIGMA shape template using the Medial Demons algorithm [1]. Registration is guided by fitting a medial model to each structure and incorporating intrinsic shape features to optimize alignment [18, 19] and two vertex-wise shape metrics are computed:

#### Radial Distance (RD)

RD quantifies local thickness by measuring the shortest Euclidean distance from each surface vertex to the medial curve of the structure. Formally, for a vertex *p* ∈ *M* on the surface and a medial curve *c*(*t*), the radial distance is defined as:

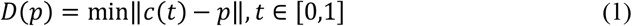

#### Tensor-Based Morphometry (TBM)

Surface-based TBM measures local deformations by quantifying the areal expansion or contraction required to align a subject’s surface mesh with a population template [20]. The differential map between the tangent spaces of the two surfaces replaces the Jacobian:

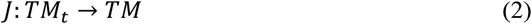

This map defines a linear transformation *J* from the tangent space of the template *M*_*t*_ to that of the subject’s surface *M* capturing local shape distortions in a surface-intrinsic manner. While full tensor analysis using Log-Euclidean metrics on symmetric positive-definite (SPD) matrices is possible [21], it is less interpretable. Therefore, we focus on the Jacobian determinant, which represents the local surface dilation ratio between the subject and the template. This measure reflects the degree of stretching or contraction required to map a small patch on the template surface to its corresponding patch on the subject. To facilitate statistical modeling, we applied a logarithmic transformation to the Jacobian determinant, yielding a final TBM feature, log-Jac, which is approximately Gaussian distributed and captures localized volumetric differences across the surface.

All image segmentation and subcortical shape models were visually inspected following ENIGMA-standardized visual quality control procedures (http://enigma.ini.usc.edu/protocols/imaging-protocols/). Approximately 96% of the segmentations passed quality control, and segmentation failures were withheld from subsequent analyses.

### 2.3 Multisite Data Harmonization

Our toolkit provides widely used data harmonization techniques when combining multisite neuroimaging data. ComBat [22] and ComBat-GAM [23] are two popular batch correction techniques used to adjust for site-related variability (e.g., scanner, sequence, field strength) while preserving, as much as possible, biologically meaningful variance across samples. The traditional linear ComBat models site effects using an empirical Bayes framework while attempting to preserve variance associated with biological covariates such as diagnosis, age, and sex. We used a Python-based adaptation of the original R implementation, available at https://github.com/Jfortin1/neuroCombat/.

ComBat-GAM extends this framework by incorporating nonlinear covariate effects using generalized additive models (GAMs). In our analysis, age was modeled as a smooth term, while diagnosis and sex were treated as categorical variables. Default empirical Bayes settings were used for site effect estimation, and no custom constraints were applied to the smoothing terms. ComBat-GAM harmonization was included in adapting Python scripts available at https://github.com/rpomponio/neuroHarmonize.

### 2.4 Mass-Univariate Statistical Models

Our toolkit supports mass univariate statistical analysis to identify vertex-wise associations between brain morphometry and clinical or demographic variables of interest across all subcortical structures. This approach fits a general linear model (GLM) or linear mixed-effects model independently at each vertex, enabling spatially detailed statistical inference across the brain, as in our prior work [10–13]. Mixed models can also be used to fit random effects (site, family structure, etc.)

To more efficiently perform vertex-wise mixed-effects modeling on large-scale neuroimaging consortium and biobank datasets, we integrated **FEMA (Fast and Efficient Mixed-Effects Algorithm)** [14] into our toolkit. FEMA uses sparse matrix algebra and parallel computation to reduce runtime and memory usage, making it suitable for high-dimensional analyses. Here, FEMA is provided as a callable module that can be applied to subcortical shape data, offering a fast and reproducible pipeline for statistical analysis compared to traditional linear models.

To assess computational efficiency, we compared the time required to fit a linear mixed-effects model using our FEMA framework versus MATLAB’s built-in *fitlme* function [24]. Both methods were applied to the same model: measure ∼ diagnosis + age + sex + ICV + (1|Site) across 14 subcortical structures comprising a total of 27,120 vertices for 3,373 participants. All results were corrected for multiple comparisons (False Discovery Rate (FDR) [25] *q*<0.05).

### 2.5 Machine Learning Analysis

Our toolkit includes a logistic regression model with structured sparsity, incorporating an L1-norm penalty to promote sparsity and a Total Variation (TV) norm to enforce spatial smoothness in vertex-wise coefficients [16]. This joint regularization approach, commonly referred to as Logit-TVL1, can enhance model interpretability and has been widely applied in neuroimaging, including diffusion MRI studies of neurological disease [15], and recent extensions to mesh-based regression and spectral analysis [26]. Unlike mass-univariate statistical approaches, which assess each vertex independently, Logit-TVL1 models the joint contribution of spatially coherent regions, capturing multivariate structure in the data.

To test the framework, we applied this model to the example task of BD versus HC diagnostic classification as well as treatment at time of scan prediction using vertex-wise subcortical thickness features extracted from 14 subcortical regions across both hemispheres. We used ComBat-GAM harmonized features to mitigate potential site effects. We performed a grid search with five-fold cross-validation to search for optimal hyperparameters, L1 and TV penalties. Model performance was evaluated using ROC-AUC score and balanced accuracy.

### 2.6 Results Visualizations

The toolkit provides multiple visualization styles to texture modeling results onto the ENIGMA subcortical shape template (**Fig. 2**): In the **original view**, brain structures are displayed in their native anatomic spatial configuration. The **unpacked view** separates the subcortical structures to reveal dorsal/ventral and lateral/medial surfaces while maintaining a loose anatomical correspondence. The **interactive view** enables dynamic 3D exploration of the subcortical brain shapes and vertex-wise coefficients in a web browser, so that users can zoom, rotate, and inspect structures and statistical analysis results with greater flexibility. The example of the interactive view is available at https://yim-igc.github.io/ENIGMA_Shape_Visualization/.

**Fig. 2.**
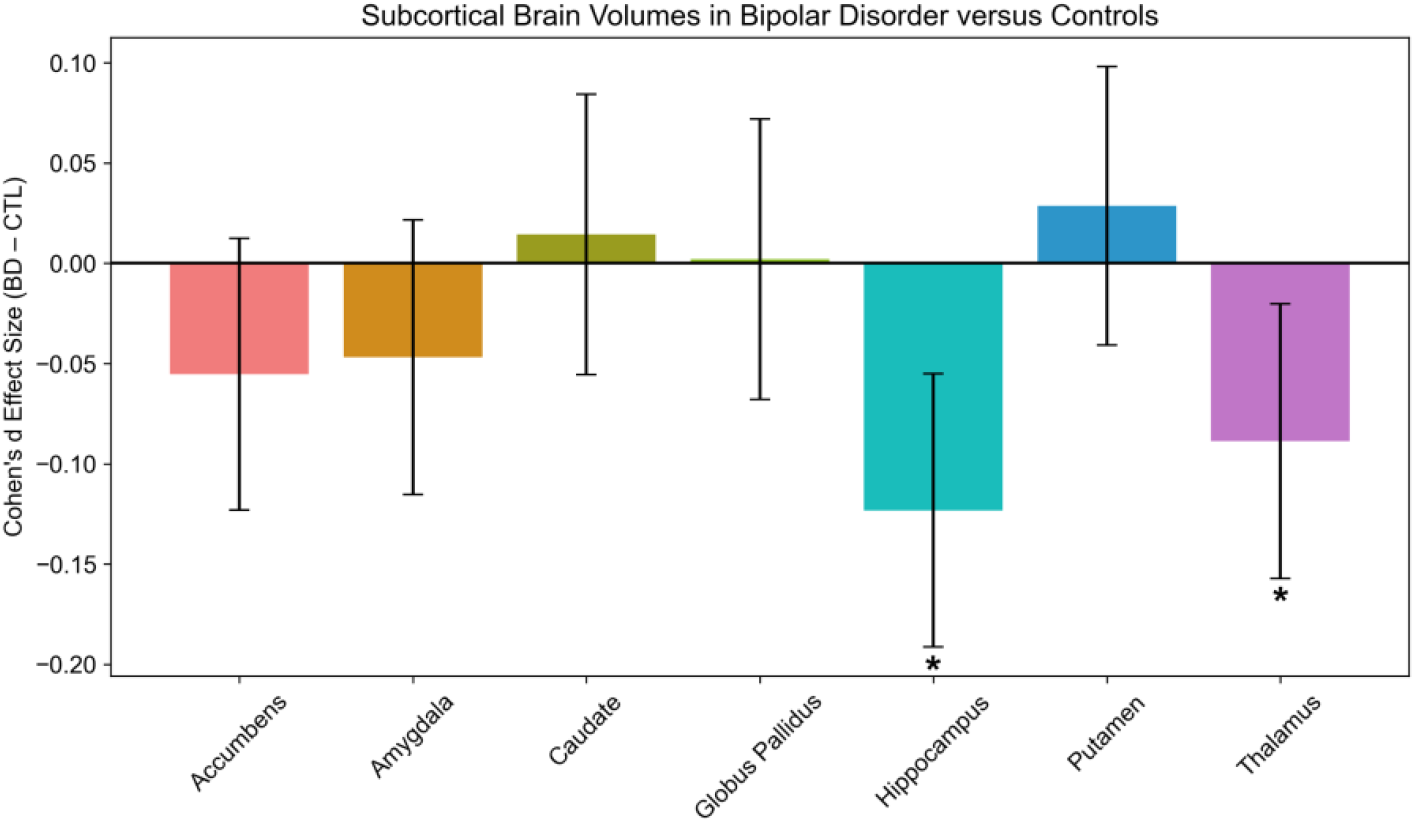
Cohen’s d effect sizes from gross volumetric analysis of BD versus HC using mixed-effect models, accounting for site as a random effect and adjusting for age, sex and ICV in 21 cohorts from ENIGMA-BD. Effect sizes were considered significant (marked with *) if they passed FDR correction for multiple comparisons.

## 3 Experimental Results

### 3.1 Gross volume and subcortical shape alterations in BD

Gross volumetric analysis showed that those with BD had smaller hippocampal (Cohen’s *d* = −0.12) and thalamic volumes (*d* = −0.08) compared to HC (**Fig. 2**), in line with our prior work [6].

Subcortical shape analysis (**Fig. 3**) revealed complex differences between BD and HC that were not detected in the gross volumetric analysis. Hippocampal and thalamic shapes mainly showed locally thinner and smaller surface area in those with BD compared to HC. The putamen and caudate displayed more complex alterations, including local regions with greater surface area as well as thinner radial distance in BD. Interactive visualizations of these preliminary findings are available at https://yim-igc.github.io/ENIGMA_BD_Shape_LME/, https://yim-igc.github.io/ENIGMA_BD_Shape_ComBat/, https://yim-igc.github.io/ENIGMA_BD_Shape_ComBat-GAM/.

**Fig. 3.**
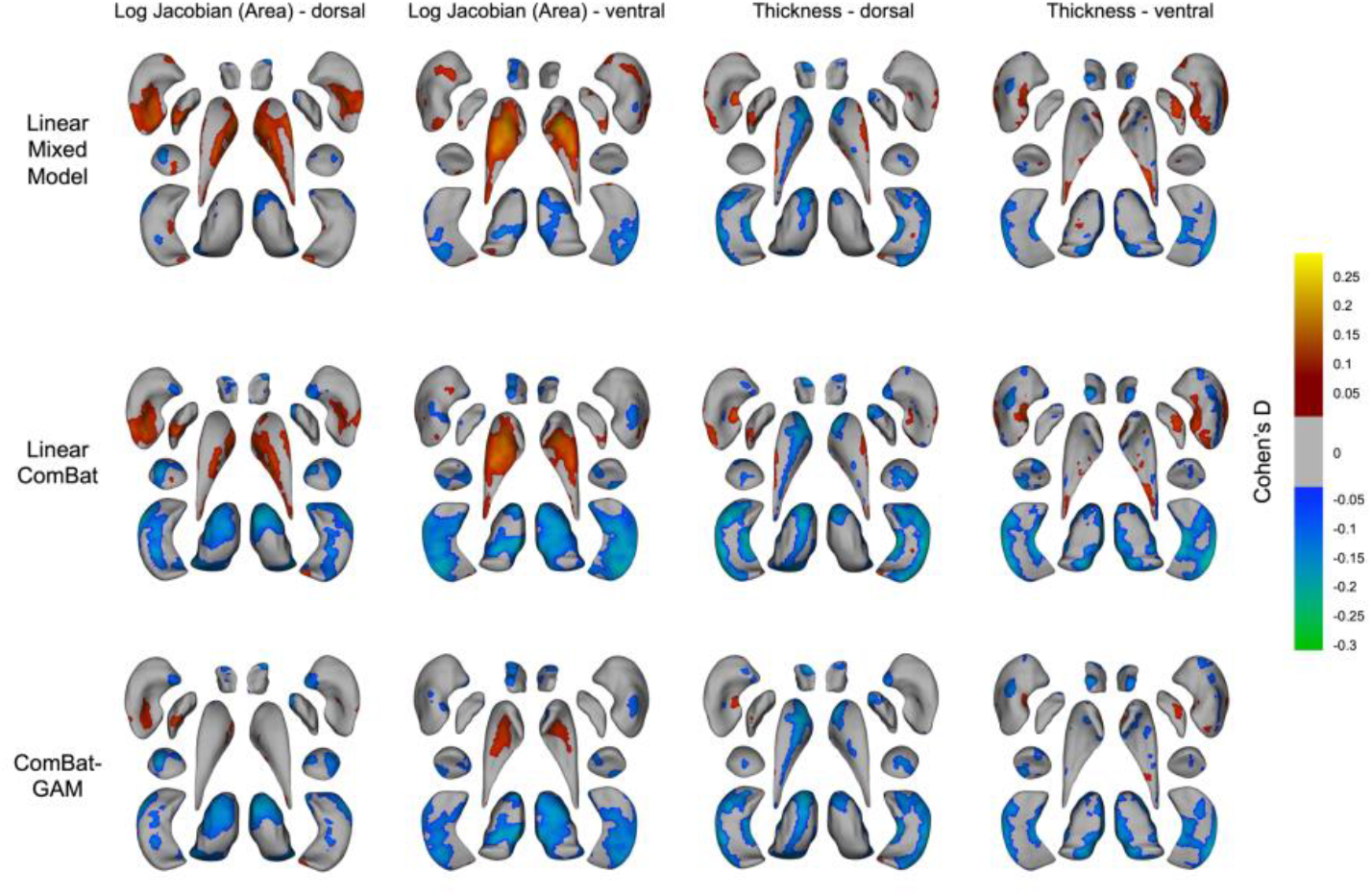
Cohen’s D maps from shape analysis using mixed-effects models that account for site as a random effect and adjusted for age, sex and intracranial volume (ICV). Gray areas indicate regions without statistically significant difference after false discovery rate (FDR) multiple comparison correction.

Application of different site harmonization techniques (site-as-random effect linear model and ComBat versions) resulted in similar patterns of BD vs. HC shape patterns, though the extent of significant group differences after correction for multiple comparisons varied to some degree between methods (**Fig. 3**). Mixed models revealed significant differences in radial distance (RD) at 33% of the vertices and in TBM at 34% of the vertices. Linear ComBat-corrected data analysis showed significant differences in RD at 42% of the vertices and in TBM at 47% of the vertices. For ComBat-GAM–corrected data, significant differences were observed in RD at 35% of the vertices and in TBM at 37% of the vertices.

### 3.2 Computational Time of Statistical Analysis

#### Model acceleration with FEMA

FEMA reduced computation time 16-fold compared to the traditional linear approach (*fitlme*) as shown in **Table 2**.

**Table 2.** Computation time for FEMA and *fitlme* in MATLAB for one model computed across all 14 structures (27,120 vertices in total) for 3,373 participants. Model: measure ∼ diagnosis + age + sex + ICV + (1|Site).

|  | FEMA | *fitlme* |
| --- | --- | --- |
| Computation time (sec.) | 60.74 | 992.41 |

### 3.3 Multivariate Analysis

Structured sparse logistic regression (Logit-TVL1) on a diagnostic classification (BD vs HC) using vertex-wise local thickness features achieved a ROC-AUC of **0.56** and a balanced accuracy of **0.56**. The predictive model for lithium treatment at time of scan achieved a ROC-AUC of **0.62** and a balanced accuracy of **0.62**, while the model for antiepileptic treatment yielded a ROC-AUC of **0.67** and a balanced accuracy of **0.67**.

## 4 Discussion

In this study, we provide a scalable toolkit for large-scale multisite consortium studies to perform 3D surface-based brain geometry modeling and demonstrate an example analysis using a large, multisite sample from the ENIGMA Bipolar Disorder Working Group. By integrating robust mass-univariate modeling, harmonization techniques, and shape-based AI methods, our toolkit supports a wide-range of analysis applications including mapping diagnostic group differences and associations with clinical and demographic variables such as treatment status.

Integration of FEMA for accelerated statistical modeling addresses a key computational bottleneck in large-scale shape-based analyses. Traditional vertex-wise mixed-effects modeling is often limited by prohibitively long runtimes and high memory demands, particularly in consortium or biobank settings including thousands of participants and tens of thousands of surface features. In our benchmark comparison, FEMA achieved a 16-fold reduction in computation time relative to MATLAB’s standard *fitlme* function. This acceleration allows researchers to conduct high-dimensional, spatially resolved modeling across large datasets in a more replicable, scalable manner.

Management of potential site effects is essential for large-scale neuroimaging studies. Our integration of ComBat and ComBat-GAM into the pipeline enables robust adjustment for these confounds while preserving meaningful biological variance. In our analyses, both methods produced consistent spatial patterns of BD versus HC group differences, though the extent of statistically significant findings varied slightly across approaches. These results highlight how site-based harmonization strategies can influence downstream analyses.

The integration of structured sparse logistic regression (Logit-TVL1) into our pipeline provides a powerful framework for machine learning analysis of subcortical shape features. In contrast to conventional univariate models that evaluate each vertex independently, Logit-TVL1 models spatially distributed patterns by combining sparsity and spatial smoothness constraints. This is particularly valuable in the context of psychiatric disorders, where anatomical alterations may be subtle, diffuse, and highly variable across individuals. The modest diagnostic (BD vs. HC) classification performance, and predictive models for treatment with lithium and antiepileptic use at time of scan were consistent with our prior gross volumetric [6], cortical [3], hippocampal subfields analysis [27].

Limitations of the present work include the following: First, the current implementation of the toolkit primarily supports subcortical shape data. Work is underway to expand functionality to support cortical surface shape data. Second, our current machine learning modeling framework focuses on Logit-TVL1. We are working to incorporate other explainable AI models such as spherical CNNs [28] or mesh-based vision transformers [29] which may improve task performance and provide deeper insight into more clinically-relevant brain alterations in BD and other disorders. Future work is also underway to combine the surface-based brain features with data from other biological scales such as polygenic risk, gene expression cytoarchitecture and neurotransmitter topography [30, 31].

## Requirements and Code availability

This toolkit is compatible with Python 3.8.x in major operating systems such as Mac OS X, Windows, and Linux. Our toolkit requires independent installation of the following Python packages: *pandas* (v2.0.x), *numpy* (v1.24.x), and *scikit-learn* (v1.3.x). Development is ongoing to incorporate additional modules that accommodate diverse types of shape data and AI models. All code is available upon request. Once it is fully validated, it will be made publicly accessible on the ENIGMA GitHub repository (https://github.com/ENIGMA-git).

## Acknowledgments

This work was supported by NIH grants R01 MH129742, AG081571 and R21 MH139001, R56 AG058854, and the Baszucki Brain Research Fund and the Milken Institute Center for Strategic Philanthropy grant. The BiDirect was supported by grants of the German Ministry of Research and Education (BMBF) to the University of Muenster (01ER0816 and 01ER1506). This work was funded by the German Research Foundation (DFG, grant FOR2107 DA1151/5-1, DA1151/5-2, DA1151/9-1, DA1151/10-1, DA1151/11-1 to UD; SFB/TRR 393, project grant no 521379614) and the Interdisciplinary Center for Clinical Research (IZKF) of the medical faculty of Münster (grant Dan3/022/22 to UD). The MUSC was funded by K23 AA020842. The Houston cohort was partially supported by NIMH (1R01MH085667-01A1), John S. Dunn Foundation (Houston, Texas), and Pat Rutherford Chair in Psychiatry (UTHealth Houston). The Paris cohort was funded by the French Agence Nationale Pour la Recherche. The Sydney (UNSW) study was supported by the Australian National Medical and Health Research Council (NHMRC) Program Grant 1037196 to Philip B. Mitchell, NHMRC Project Grants 1066177 and 1063960 to Janice M. Fullerton, NHMRC Investigator Grants 1176716 to Peter R. Schofield and 1177991 and Philip B. Mitchell, and NHMRC & Medical Research Futures Fund Grant 1200428 to Janice M. Fullerton. Janice M. Fullerton is the grateful recipient of the Janette Mary O’Neil Research Fellowship. Additional philanthropic support was provided by the Lansdowne Foundation, GoodTalk charity, the Gordon Pettigrew Family, Mrs Betty C. Lynch OAM (dec) and the Aberdeen Foundation. Pravesh Parekh was supported by the European Union’s Horizon 2020 research and innovation programme under the Marie Skłodowska-Curie grant 801133; Research Council of Norway grant 324252; NIH grants U24 DA041123 and U24 DA055330. Anders M. Dale is supported by NIH grants U24 DA041123, DA055330, R01 AG076838, and OT2 HL161847. Dr. Tamsyn Van Rheenen was supported by an Al and Val Rosenstrauss Fellowship from the Rebecca L Cooper Medical Research Foundation. James A. Karantonis was supported by Swinburne University/an Australian Postgraduate Award. Lisa Furlong was supported by the Australian Rotary Health and the Ian Parker Bipolar Research Fund; Brunslea Park Estate. Susan Rossell was supported by an NHMRC Senior Fellowship. The GIPSI cohort was supported by PRISMA UT and MINCIENCIAS. The COGSBD study was financially supported by the NHMRC (1060664), Henry Freeman Trust, Jack Brockhoff Foundation, University of Melbourne, Barbara Dicker Brain Sciences Foundation, Rebecca L Cooper Foundation and the Society of Mental Health Research. The authors acknowledge the facilities and scientific assistance of the National Imaging Facility, a National Collaborative Research Infrastructure Strategy (NCRIS) capability, at the Swinburne Neuroimaging Facility, Swinburne University of Technology.

